# The allometry of discontinuous gas exchange cycles in *Atta* cephalotes leaf-cutter ants

**DOI:** 10.1101/2025.11.26.690668

**Authors:** O.K. Walthaus, D. Labonte

**Affiliations:** Imperial College London; Imperial College

## Abstract

Many idle insects exhibit discontinuous gas exchange cycles (DCGs). During DCGs, CO_2_ is released in discrete bursts, followed by periods of negligible gas exchange. The standard metabolic rate (SMR) is thus determined to first order by the product between cycle frequency (f_c_) and burst volume (V_b_, SMR ≈ f_c_· V_b_). The evolutionary allometry of these parameters is well studied, but it remains unclear if their static allometry, measured in individuals of the same species, sharing the same ontogenetic stage, follows the same patterns. To address this question, we investigate the static allometry of DCGs in Atta cephalotes leaf-cutter ants workers varying by two orders of magnitude in body mass. The SMR allometry significantly exceeded the standard prediction from the nutrient supply network model, and differed from the SMR allometry observed across insects. This disproportional increase was exclusively achieved by an increase in V_b_, perhaps because f_c_ is stabilised by neural and mechanical constraints. It may be necessitated by the positive allometry of the largest muscle in Atta—the mandible closer muscle—which increases with a virtually identical allometric coefficient, providing further evidence that the principles of symmorphosis may be upheld in insects.

## Introduction

Gas exchange in insects is facilitated by the tracheal system, a network of branching tubes which open to the atmosphere through pores called spiracles (1,2), thus facilitating the delivery of oxygen (O_2_) directly to, and the removal of carbon dioxide (CO_2_) from, metabolically active tissues (1,3–5). Despite dispensing with some of the components of mammalian circulatory systems, this open system supports both a large aerobic scope and efficient gas regulation at rest (1).

Idle insects exchange gases in a variety of ways, including continuous and cyclic ventilation. By far most widely reported are discontinuous gas exchange cycles (DGCs) (6). DGCs have been observed in five insect orders across 51 species (6–8), and a large body of work has been devoted to characterising their distinct phases, their adaptive significance, physiological triggers, and evolutionary origins (1,6,8).

During DGCs, CO_2_ is released in discrete bursts (O-phase, spiracles open), followed by a period of virtually no gas exchange (C-phase, spiracles closed; for flutter phase, see SI) (1). How much gas is exchanged is then determined by the product between the cycle frequency (f_c_) and the amount of CO_2_ released per burst, the burst volume, (V_b_) (7).

As for many physiological traits, it is instructive to consider how f_c_ and V_b_ change with body size. Notably, the allometry of these critical DGC parameters remains debated: f_c_ has been reported to increase, remain stable, or decrease with body mass (7). This inconsistency may stem from challenges inherent to within-species studies: static allometry relies on individuals at the same ontogenetic stage and thus is limited in size range (7), ontogenetic allometry, in turn, can be confounded by developmental and physiological changes independent of growth (9). The result of both difficulties has been a sustained uncertainty as to whether ontogenetic or static allometry generally mirrors the evolutionary allometry across insect species (7,10). To contribute to the resolution of this question, we investigate the static allometry of DGCs in the leaf-cutter ant *Atta cephalotes* (Linnaeus, 1758).

Leaf-cutter ants are a promising model organism for at least two reasons. First, they exhibit substantial adult size variation: mature workers can vary in body mass by more than two orders of magnitude (11). Second, size-variation is accompanied by significant positive allometry of the mandible closer muscle, which accounts for as much as ∼15% of the body mass in large workers (12). Associated with this positive allometry of metabolically active tissue is a positive allometry of the standard metabolic rate (SMR) (13)—but is this achieved by increased cycling frequency, volume, or a combination of both? Because cycling frequency and volumes can, at least in principle, be linked to hypotheses about the physiological and neurological drivers of DGCs (2), determining how allometric needs are accommodated—the aim of this work—may provide a small piece to this central puzzle (2).

## Materials and Methods

### Study animals

A sub-colony of approximately 10,000 *A. cephalotes* workers was isolated from a laboratory colony, and housed in a 18⍰18⍰20 cm (L⍰W⍰H) box, connected to a foraging area and a waste box. The sub-colony was situated in a climate chamber, maintained at 25°C, 60 ±5% relative humidity, and a 12-hour light/dark cycle (Weiss Technik Fitotron, Loughborough, UK); bramble and kibbled maize were provided *ad libitum*. 22 workers, weighing between 2 and 70.1mg, were collected for respirometry measurements, such that their body masses were distributed approximately evenly in log10-space.

### Flow-through respirometry

CO_2_ emission was measured via flow through respirometry, in an environment maintained at 25 ± 0.5°C (Figure 1). Air was drawn from a reservoir (50X50X30cm), which acted as a low-pass filter for fluctuations in air consistency (14), and was chemically scrubbed of CO_2_ and H_2_O using soda lime and Drierite/Ascarite/Drierite columns (Ascarite II 20–30 mesh; Thomas Scientific, Swedesboro, New Jersey, USA, Drierite 8 mesh; Sigma-Aldrich, St. Louis, Missouri, USA) (14). Scrubbed air was drawn through the system at 50ml min^-1^ via a flow-meter and pump (SS4 Sub-Sampler, Sable systems, Nevada, USA), regulated via a calibrated mass flow control valve (0-500 SCCM Air, Alicat Scientific, Tucson, AZ, USA). A cylindrical inline chamber was used in all experiments (RC, Sable Systems, Nevada, USA, ID 2cm⍰3.8cm (Ø⍰H), internal volume = 14mL, Figure 1a); its volume was matched to the flow rate to achieve a time constant of approximately 17s—more than sufficient to record CO_2_ emission in insects (15–17). CO_2_ emission was measured downstream with a resolution of ∼0.1ppm at a frequency of 1Hz, using a LI7000 infrared CO_2_ gas analyser in differential mode (LI-COR Environmental, Lincoln, USA). The gas analyser was calibrated and spanned using certified 1000ppm CO_2_ calibration gas in balance N_2_ (BOC, certificate number: 21/005083, UK); pure N_2_ was used to calibrate to true zero (BOC, 99.99% pure nitrogen, certificate number: 21/004497, UK). Water vapour pressure, relative humidity (RH-300, Sable systems, Nevada, USA) and temperature (Type-T thermocouple probe, Sable systems, Nevada, USA) were also recorded. Raw data were relayed to Expedata v1.9.13 *via* a universal interface (UI-3, Sable Systems).

**Figure 1.**
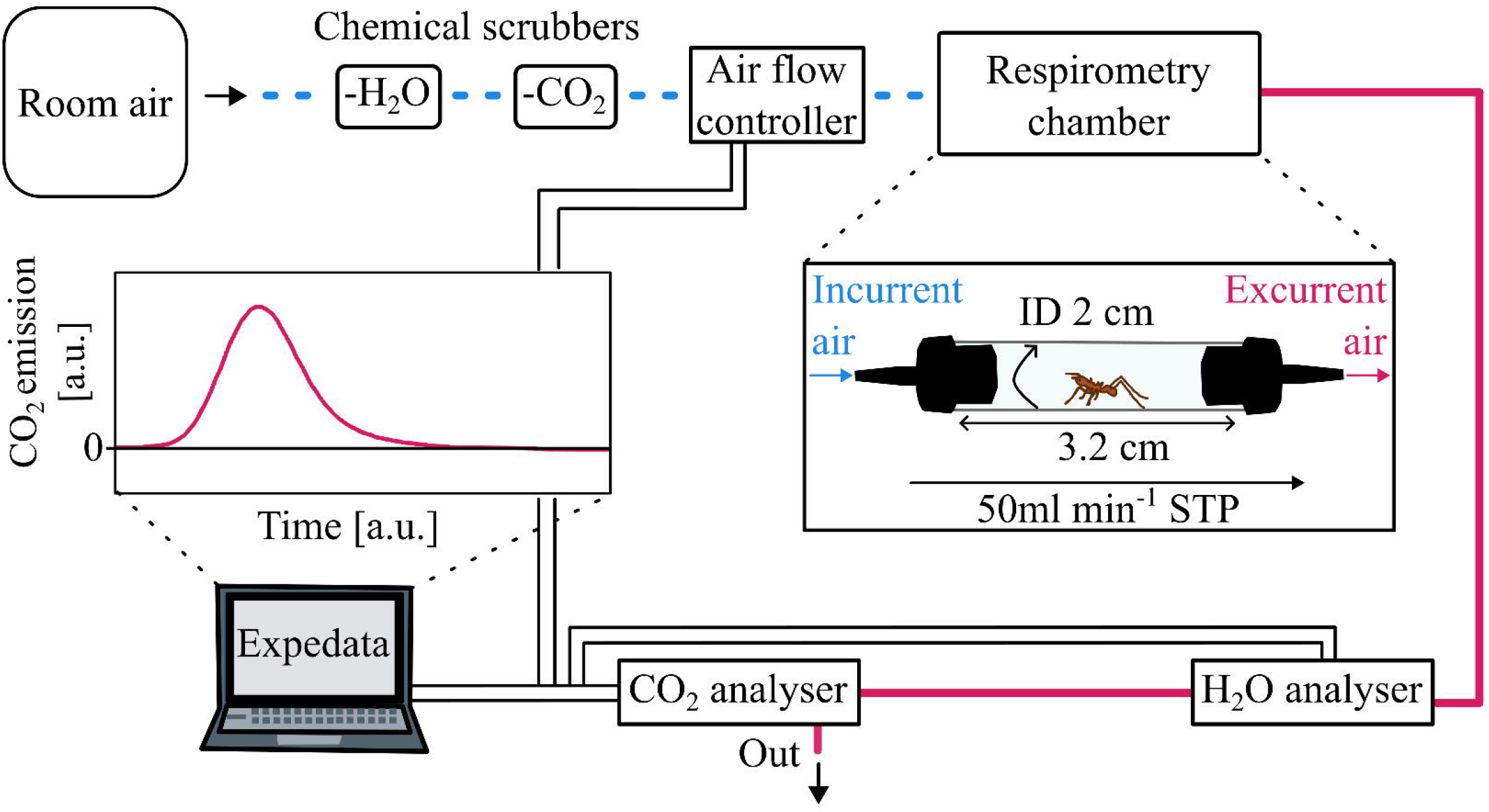
Block diagram of the flow-through respirometry setup. Blue dashed lines indicate incurrent air, pink solid lines excurrent air, and black double lines data connections. Ambient air is chemically scrubbed of carbon dioxide (-CO_2_) and water vapour (-H_2_O) by means of scrubbing columns. CO_2_ emission was measured for workers spanning 1.5 orders in body mass (2.0-70.1mg). Air was pulled through the system at 50ml min ^-1^, yielding a time constant of 17s for a chamber with a volume of 14mL. Idle individuals exhibited discontinuous gas exchange cycles (DGC): gases were exchanged during distinct bursts with a characteristic frequency, f_c_, and volume, V_b_. *Attine* ants use lipids as primary metabolite (13), defining the conversion of CO_2_ flow rates into a standard metabolic power consumption in μW.

### Experimental procedure

Idle ants were extracted from the foraging area, weighed to the nearest 0.1mg (Kern ADB 100-4 120g⍰0.001g, Kern, Germany), and placed into the respirometry chamber to acclimatise for 30-minutes (15). Each measurement began and ended with a 5-minute baselining period, taken from airflow bypassing the chamber (14). Using a central baselining unit (CBL, Sable Systems, Nevada, USA), the airstream was then switched to the active chamber, which was first flushed of CO_2_ for 5-minutes to remove accumulated gas; CO_2_ emission was then recorded for 30-min, after which the airstream was switched back.

Experiments were deemed invalid if ants (i) became active; (ii) exhibited escape or stress responses; or (iii) performed small movements that caused a clear departure from DGCs (see SI, and (15)).

### Data curation and statistical analysis

All handling of raw data—including unit conversions, marker and drift correction, baseline subtraction, automatic peak detection, and integration—was completed in Expedata v1.9.13.

The following parameters were extracted: (i) f_c_ (ii) V_b_, and (iii) SMR. f_c_ was the inverse of the average time between successive peaks (Figure 2a). V_b_ was calculated by integrating each CO_2_ burst with respect to time, (Figure 2b, see SI); a low long-term drift of 0.4ppm h^-1^ ensured that V_b_ was not significantly affected by drift (14,17). SMR was determined by multiplying f_c_ and V_b_ (Figure 2c, see SI for flutter phase).

**Figure 2.**
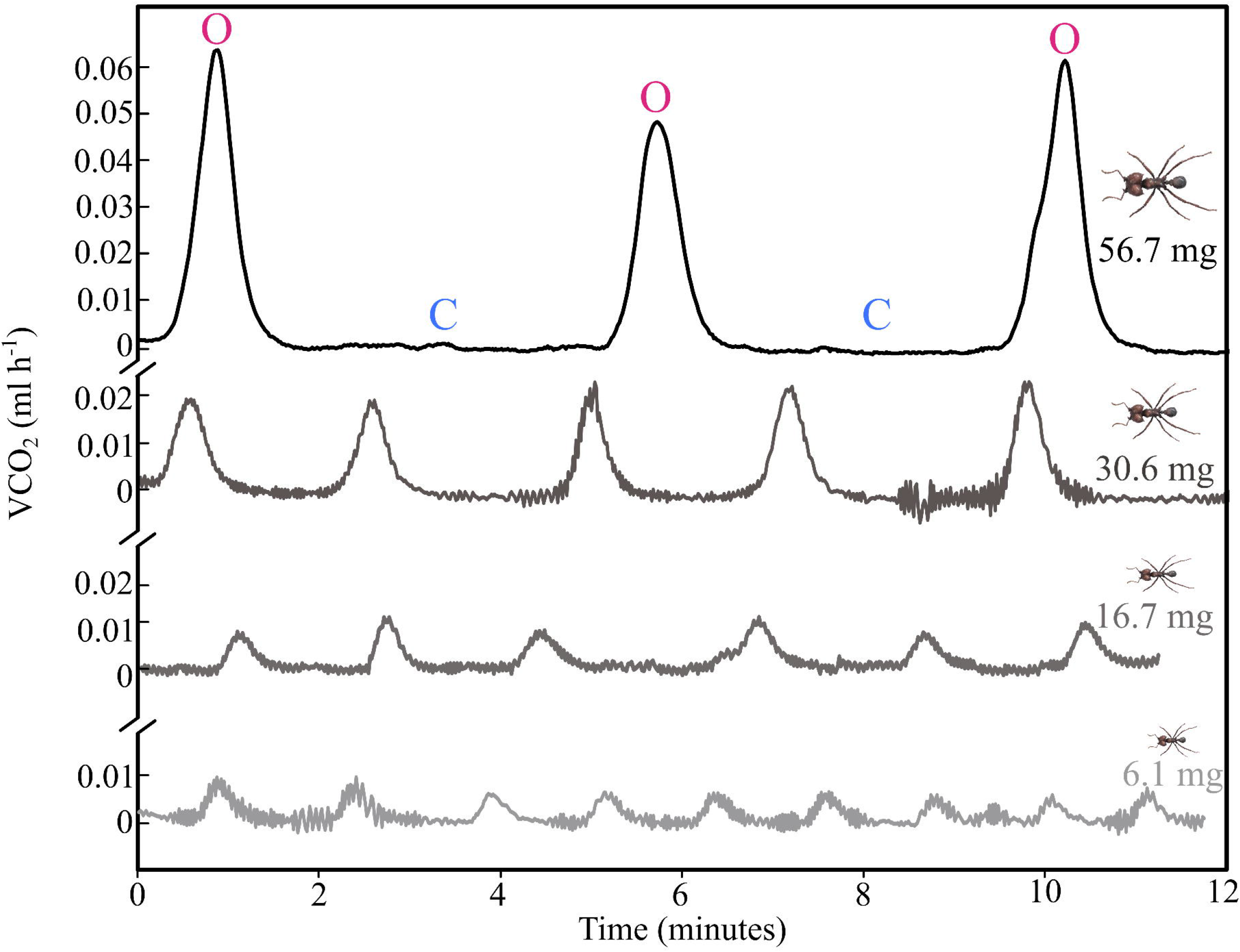
Illustrative CO_2_ emission traces for four workers of different body masses. Idle workers exhibited discontinuous gas exchange cycles (DGC), readily identifiable by distinct spiracle open (O) and prolonged closed (C) phases. The volume of CO_2_ released per burst increased with size, but the cycle frequency decreased; note well the breaks in the y-axis.

CO □emission rate was converted to metabolic power assuming lipid metabolism (RQ = 0.71) and an energy equivalent of 27.6 J mL^−1^ CO□(13). Conversions from CO_2_ emission rate to metabolic power, statistical analyses, and data curation were conducted in Rv.4.1.1. Ordinary least squares (OLS) regressions on log10-transformed data were performed to characterise allometric relationships with respect to body mass. Analysis of variance (ANOVA) was used to investigate the effect of body mass on V_b_, f_c_, and SMR.

## Results

Idle *Atta cephalotes* exhibited DGCs, readily identifiable from prolonged periods of spiracle closure (C), and distinct open phases (O) (6,18) (Figure 2).

The cycle frequency, f_c_, decreased significantly with body mass, (ANOVA: F_1,20_ = 75.3, p < 0.001), dropping from a maximum of 10.2mHz to a minimum of 3.5mHz (Figure 3a). This allometric pattern is satisfactorily described as f_c_ = 13.77m^-0.29^ (R^2^ = 0.79), where body mass, *m*, is in mg and f_c_, is in mHz (95% CIs: [10.96, 17.38] and [-0.36, -0.22] for pre-factor and scaling exponent, respectively).

**Figure 3.**
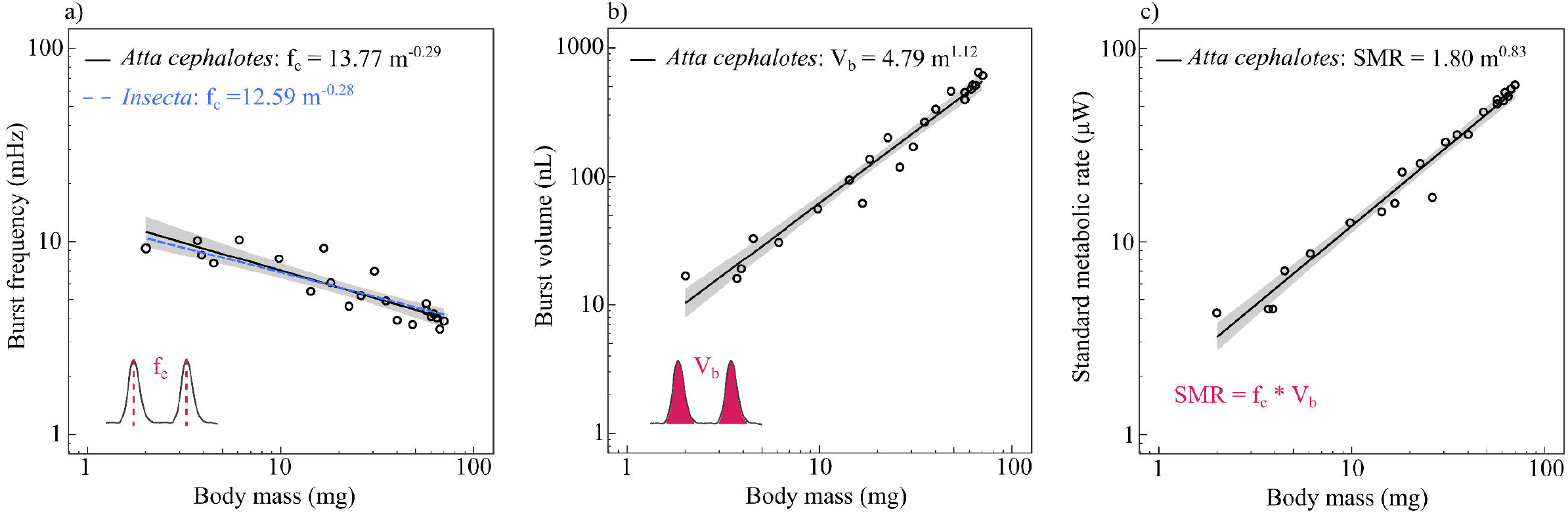
To characterise the allometry of the discontinuous gas exchange cycles, (i) cycle frequency, (ii) burst volume, and (iii) standard metabolic rate were extracted from CO_2_ emission recordings of *Atta cephalotes* workers varying by 1.5 orders of magnitude in body mass (2-70.1mg). **a)** Cycle frequency (f_c_) decreased significantly with body mass, f_c_ ∼ m^-0.29^, and ranged from 3.5–10.2mHz. This slope is compared to the result of a meta-analysis including 80 insect species across 14 orders (blue dashed line, phylogenetic generalised least-squares with coefficient for gas exchange type) (7). **b)** Burst volume (V_b_) increased significantly with body mass: the largest worker released about 40 times more CO_2_ than the smallest worker, V_b_ ∼ m^1.12^. **c)** The standard metabolic rate (SMR) is equal to the product of f_c_ and V_b_, SMR ∼ m^0.83^. All data are plotted in log-log space; lines and shaded regions indicate the allometric relationships and their confidence intervals as estimated via ordinary least-squares regressions.

The amount of CO_2_ released during the O-phase—the burst volume V_b_—increased significantly with body mass (ANOVA: F_1,20_ = 536.1, p < 0.01); the allometric relationship is V_b_ = 4.79m^1.12^ (R^2^ = 0.96), where body mass, *m*, is in mg and the volume, V_b_, is in nL (95% CIs: [3.39, 6.46] and [1.02, 1.22] for pre-factor and scaling exponent, respectively). Thus, the largest worker released about 40 times more CO_2_ per burst than the smallest worker (Figure 3b).

The size-specific variation in burst volume and cycle frequency combined to a substantial increase of the SMR (ANOVA: F _1,20_ = 727.4, p<0.01), SMR = 1.80m^0.83^ (R^2^ = 0.97), where body mass *m* is in mg and SMR is in μW (95% CIs: [1.46, 2.21] and [0.76, 0.89] for pre-factor and scaling exponent, respectively). The smallest worker consumed about 4μW when idle; the largest worker used about 25 times more (60µW, Figure 3c).

## Discussion

Be they active or at rest, insects need to ensure a consistent supply and timely removal of oxygen and toxic carbon dioxide. In many idle insects, both requirements are met by discontinuous gas exchange cycles (DGCs) (1,6–8) DGCs are characterised by a cycle frequency, f_c_, and a burst volume, V_b_, which jointly define the standard metabolic rate, SMR ≈ f_c_ · V_b_. The evolutionary allometry of DGC parameters is well studied, (6–8), but their static allometry has remained less clear, partially because the limited size ranges of within-species studies increase statistical uncertainty. To circumvent this difficulty, we measured DGCs in mature leaf-cutter ant workers spanning two orders of magnitude in body mass.

Consistent with earlier work on leaf-cutters (13), the SMR of *A*.*cephalotes* grew more rapidly than predicted by the null hypothesis from the nutrient supply network model (NSNM), SMR_NSN_ ∝ m^0.75^ (19). A more rapid SMR growth is a common observation in static or ontogenetic allometries in insects, but contrasts with the evolutionary SMR allometry, which follows the null hypothesis closely (9,20). The positive static allometry of the SMR raises four questions, which we now address.

First, a positive SMR allometry can be achieved by more rapid growth of V_b_, a shallower drop of f_c_, or a combination of both. Which strategy is implemented? Combining the NSNM with isogeometry yields f_c_ ∝ SMR_NSN_ · V_b_^-1^ = m^0.75-1^ = m^-0.25^, a prediction that is statistically indistinguishable from the static allometry observed here, and consistent with the evolutionary allometry of f_c_ in both vertebrates and invertebrates, despite striking differences in respiratory systems (6,7,20,21). Thus, the positive allometry in SMR appears to be exclusively achieved by a positive allometry in V_b_, V_b_ ∝ m^1.12^.

Second, it is not immediately obvious why a positive SMR allometry may be preferentially implemented *via* a disproportionate increase in V_b_. What underpins the seeming invariance of the size-specific cycling frequency? DGCs consistently correlate with reduced brain activity (2). In normoxic conditions, the opening and closing of spiracles is instead controlled by central pattern generators (CPGs) (2), which respond to CO_2_ and O_2_ partial pressure (1,2): spiracle opening is triggered by an accumulation of CO_2_ to near-critical levels (8,18), enabling rapid expulsion down a steep partial pressure gradient (22). Although gas exchange can also be modulated by active neurological control—as observed in hypercapnic locusts or cockroaches (2)—this (i) will likely reduce gas exchange efficiency due to smaller partial pressure gradients, and (ii) be more costly, as it requires the use of complex neural circuitry (2); it is thus perhaps the inferior strategy.

Third, a size-specific increase in V_b_ despite equal size-specific cycling times will either require active ventilation, typically observed in larger insects (3,23), or more efficient removal of CO_2_ by other means, e.g., by facilitating a more rapid build-up of pressure gradients. How may this be accomplished? The volume of the insect tracheal system tends to scale with positive allometry (V_tracheal_ ∼ m^1.02–1.24^) (3,24–27), and thus can likely accommodate larger burst volumes. In ants, the branching of the tracheal network breaks with Da Vinci’s rule obeyed in large mammalian vessels (19), and instead seems to follow Nunome’s pattern (24): the summed area of child branches becomes progressively smaller, not larger. This branching pattern increases the maximal CO_2_ flux, i.e., it facilitates outward release of CO_2_ (24); it may thus help maintain efficient gas exchanges despite an increased size-specific burst volume (24).

Fourth, the positive allometry of tracheal volume, necessary to accommodate larger V_b_, comes at a cost: it decreases the space available for other tissues. So, what makes it necessary? A positive allometry of SMR is common in within-species studies in insects (10,20), but its mechanistic origin remains contested. One candidate theory states that the SMR will be elevated above the level predicted by the NSNM if growth is dominantly achieved through increases in cell number as opposed to cell size (20). In closely related *A*.*vollenweideri*, the volume, V_m_, of the mandible closer muscle grows predominantly through the addition of new fibres rather than by fibre hypertrophy (12), providing some support for the cell size model. Our results also provide a further piece of evidence that the principles of symmorphosis—that the respiratory system meets functional requirements with minimal excess capacity (28)—may be observed in insects. More detailed work in the locust reported a congruence between the mitochondrial volume of major muscle groups and the tracheal volume, which both grew isometrically (9); here we provided initial evidence that such matching may be sustained even when muscle volume departs from isometry: the mandible closer muscle grows as V_m_ ∝ m^1.15^ (12), tantalisingly close the allometry of the burst volume, V_b_ ∝ m^1.12^. Curiously, the scaling of the SMR, SMR ∼ m^0.83^, mirrors almost exactly the scaling of the physiological cross-sectional area of the mandible closer muscle A_phys_ ∼ m^0.84^ (12).

Although this may appear superficially consistent with force-cost theories that assign a dominant role to muscle force in the determination of metabolic costs (29,30), this similarity should, at this point, not be over-interpreted; instead, we see it merely as providing a motivation for follow-up work that seeks to investigate the relation between allometries of major muscle groups and the allometry of DGCs in more detail.

