## Supplementary material for "The allometry of discontinuous gas exchange cycles in *Atta* cephalotes leaf-cutter ants"

O.K. Walthaus and D. Labonte

**Supplementary information**

**Conversion from CO_2_ in ppm to ml.s^-1^**

CO_2_ emission rate was calculated by dividing measured ppm CO_2_ by 1 million to obtain to a fraction. This fraction was then multiplied by the airflow rate (in mL·min^-1^) and divided by 60 to obtain a volume flow rate in mL·s^-1^ (1).

**Burst onset detection to calculate V_b_**

CO_2_ burst onset was calculated using a derivative-based threshold method. The raw CO_2_ traces were first smoothed using a third order Savitzky Golay filter with a 10-point window to reduce high-frequency noise while preserving the overall shape of the bursts. We then took the first derivative of the smoothed CO_2_ trace, ${dC}/{dt}$, calculated as the difference between consecutive points. A 30 second baseline period was taken to represent the baseline slope, $\mu_{base}$, with its standard deviation given as $\sigma_{base}$. Burst onset, $B_{onset}$, was given as the first point where ${dC}/{dt}$, exceeded $\mu_{base}+4\sigma_{base}$ for at least 5 consecutive seconds. Burst termination, $B_{end}$, was defined as the point where ${dC}/{dt}$, returned within $\pm\sigma_{base}$ following the burst peak, a more lenient $\sigma_{base}$ criterion was used for burst termination to ensure the full extent of the CO_2_ peak was captured. Burst CO_2_ volume was calculated by integrating under the peak between $B_{onset}$ and $B_{end}$. Average burst volume was calculated across all peaks in the measurement window and rounded to the nearest nL. A derivative based approach is recommended for burst onset detection (1) and provides an objective, reproducible method for quantifying burst volumes while excluding flutter contributions.

**\subsection{Comparing total SMR between full trace integration and cycle frequency*volume}**

Discontinuous gas exchange in insects comprises three phases of CO_2_ emission, which can be crudely described as: an open (O) phase, during which spiracles remain fully open and most CO_2_ is released (2,3); a flutter (F) phase, characterised by rapid spiracle movements and the release of about 5–15% of total CO_2_ (2,3); and a closed (C) phase, when CO_2_ release is minimal (2). These phases are often inferred from CO_2_ traces, but without direct spiracle observation and pressure measurements, distinguishing them with certainty, particularly the F phase, is difficult (2,3). As we exclude the flutter phase from the burst volume analysis we here want to account for its contribution to total SMR to confirm its exclusion *a priori* as to not neglect it entirely. In this study, we define the flutter phase as any interburst period between open phases in which CO_2_ release is above zero, indicating that spiracles are not fully closed. These periods are often visible in CO_2_ traces (SI Figure 1). We then compare standard metabolic rate (SMR) estimates derived from full-trace integration (including F phase) with those calculated from frequency * volume (neglecting F phase).

The difference between both methods provides a proxy for the contribution of the flutter phase to the SMR. The SMR estimate obtained from integration was, on average, 5% higher than the burst volume-frequency estimate, but never exceeded it by more than 10% (SI Figure 2). Because the contribution of the flutter phase is consistently small, we here simply estimate the SMR from burst-volume and frequency (but see SI Figure 2).

**SI Figure 1:**

While CO_2_ traces alone cannot fully determine the flutter phase, clear examples of flutter can be observed in the traces. An example raw trace from a quiescent 30.6 mg worker shows clear flutter (green), open (pink) and closed (blue) phases for one cycle of CO_2_ release.

**SI Figure 2:**

To discern the contribution of the flutter phase to total SMR we independently assessed SMR through full trace integration (including flutter phase contribution, grey data) and through cycle frequency * burst volume (neglecting flutter phase contributions, pink data). Both methods of analysis yielded SMR scaling approximately as $SMR \sim m^{0.82-0.83}$ with the full trace integration being on average 5% higher indicating the flutter phase contributes approximately 5% to total CO_2_ production.

**Activity affects discontinuous gas exchange cycles**

**SI Figure 3:**

Even minor movements small movements (e.g., leg tapping, minor head rotations) can have a significant effect on the discontinuous gas exchange pattern of idle workers. The example here shows the metabolic trace of a 43.0 mg worker that slowly rotated during the measurement. Note the elevated trace that does not return to zero, suggesting the spiracles never fully close. In addition, burst peaks are irregular.

**Mass scaling exponents across *Atta* species**

**SI Table 1:** Scaling of standard metabolic rate in *Atta* ant species. ***** Denotes values obtained from the literature (4,5). Size range refers to the orders of magnitude difference between the largest and smallest individuals measured. Subscript values represent CI

| Species | n | Size range | Mass scaling exponent |
| --- | --- | --- | --- |
| *Atta cephalotes* | 22 | 1.52 | _0.78_ 0.83 _0.88_ |
| *Atta vollenweideri* | 8 | 0.95 | _0.80_ 0.88 _0.96_ |
| *Atta columbica******** | 27 | 0.97 | _0.44_ 0.64 _0.84_ |
| *Atta sexdens******** | 18 | 0.91 | _0.87_ 0.89 _0.92_ |
