## Supplementary figures and images for "The allometry of discontinuous gas exchange cycles in *Atta* cephalotes leaf-cutter ants"

### SI_figure_1

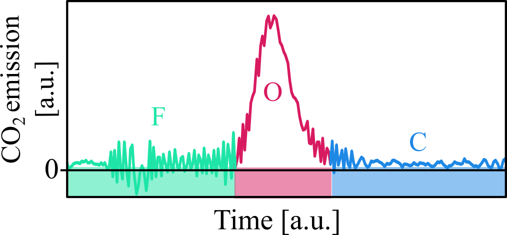

### SI_Figure_2

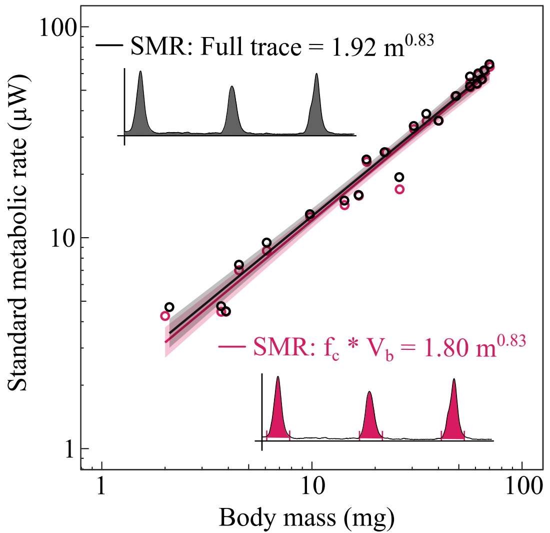

### SI_Figure_3

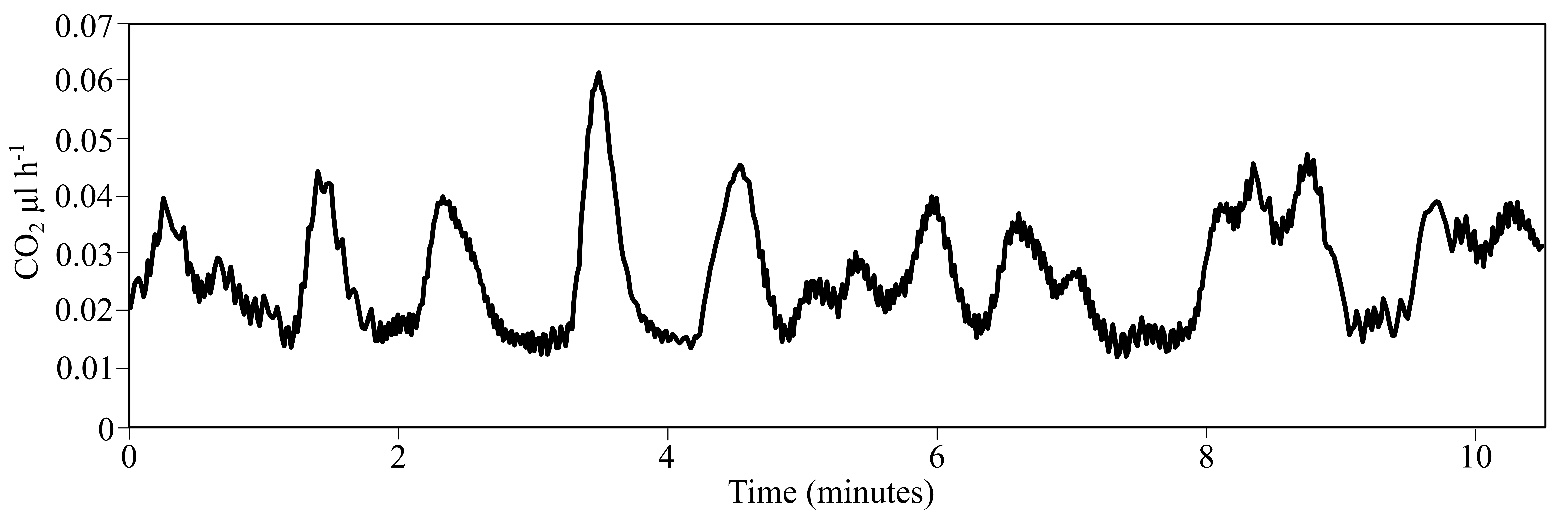
